# Distinct structural motifs are necessary for targeting and import of Tim17 in *Trypanosoma brucei* mitochondrion

**DOI:** 10.1101/2023.07.07.548172

**Authors:** Chauncey Darden, Joseph Donkor, Olga Korolkova, Muhammad Younas Khan Barozai, Minu Chaudhuri

## Abstract

Nuclear-encoded mitochondrial proteins are correctly translocated to their proper sub-mitochondrial destination using location specific mitochondrial targeting signals (MTSs) and via multi-protein import machineries (translocases) in the outer and inner mitochondrial membranes (TOM and TIMs, respectively). However, MTSs of multi-pass Tims are less defined. Here we report the characterization of the MTSs of *Trypanosoma brucei* Tim17 (TbTim17), an essential component of the most divergent TIM complex. TbTim17 possesses a characteristic secondary structure including four predicted transmembrane (TM) domains in the center with hydrophilic N- and C-termini. After examining mitochondrial localization of various deletion and site-directed mutants of TbTim17 in *T. brucei* using subcellular fractionation and confocal microscopy we located at least two internal signals, 1) within TM1 (31-50 AAs) and 2) TM4 + Loop 3 (120-136 AAs). Both signals are required for proper targeting and integration of TbTim17 in the membrane. Furthermore, a positively charged residue (K^122^) is critical for mitochondrial localization of TbTim17. This is the first report of characterizing the internal mitochondrial targeting signals (ITS) for a multipass inner membrane protein in a divergent eukaryote, like *T. brucei*.

**Summary:** Internal targeting signals within the TM1, TM4 with Loop 3, and residue K122 are required collectively for import and integration of TbTim17 in the *T. brucei* mitochondrion. This information could be utilized to block parasite growth.

## 1. Introduction

Mitochondria are essential organelles in eukaryotes. This is not only because mitochondria produce 80% of cellular ATP, but also for their roles in various cellular functions, e.g., calcium homeostasis, programmed cell death, antiviral immunity, and cell signaling (Walker et al., 2014; Weinberg et al., 2015; Mehta et al., 2017; Eisner et al., 2018; Calvo-Rodriguez and Bacskai, 2020). Mitochondrial defects cause many human diseases particularly several devastating neurodegenerative disorders including Alzheimer’s disease, Parkinson’s disease and others (Nunnari and Suomalainen, 2012; Bogorodskiy, et al., 2021; Ciccarelli et al., 2023; Mambro et al., 2023; Rehman et al., 2023). In order for mitochondria to perform such diverse functions, close to 1,000 nuclear-encoded proteins have to be imported into the mitochondria from the cytosol (Ozaowa et al., 2003; Schulte et al., 2023). Mitochondria possess an intricate machinery to import these nuclear-encoded proteins and to sort them into proper sub-mitochondrial destinations (Schmidt et al., 2010; Neupert 1997).

Mitochondrial import machinery has long been studied in yeast, human, and plants (Murcha et al., 2003; Schmidt et al., 2010; Suzuki et al., 2000; Ghifari et al., 2018) In these organisms, the mitochondrial outer membrane (MOM) possesses one translocase, the TOM complex that imports almost all nuclear-encoded mitochondrial proteins (Drwesh and Rapaport, 2020). Whereas, the mitochondrial inner membrane (MIM) has two translocases, TIM23 and TIM22 (Schmidt et al., 2010; Neupert 1997). Likewise, TOM and TIMs are multi-subunit protein complexes. Mitochondrial matrix proteins with N-terminal targeting signal (MTS), also referred to as the presequence, are translocated from the TOM to the TIM23 complex (Bomer et al., 1996; Neupert and Brunner, 2002). Whereas, polytopic mitochondrial inner membrane proteins are translocated via the TIM22 complex (Sirrenberg et al., 1996; Koehler et al., 1998; Kolli et al., 2018). These proteins are chaperoned from the TOM to the TIM22 complex via small Tim complexes in the IMS (Gebert et al., 2008; Baker et al., 2009). The major channel-forming unit of the TIM23 and TIM22 complexes are Tim23 and Tim22, respectively (Truscott et al., 2001; Rehling et al., 2003). Tim17 interacts with Tim23 and acts as a structural component of the TIM23 translocase and regulates its activity (Martinez-Caballero et al., 2007). Mitochondrial membrane potential is required for protein translocation through either the TIM23 or TIM22 complexes (Schmidt et al., 2010; Neupert 1997). In spite of elaborate discoveries of mitochondrial protein import machinery in fungi and mammals, protein complexes involved in similar functions in early divergent eukaryotes have just began to emerge (Chaudhuri et al., 2020; Schneider 2022).

*Trypanosoma brucei* is a parasitic protozoan that belongs to a group of very early divergent eukaryotes. *T. brucei* causes a fatal disease in humans known as African Sleeping Sickness and a similar disease in livestock, called Wasting Syndrome or Nagana (Sternberg and Maclean, 2010; Kasozi et al., 2022). One of the several unique characteristics of *T. brucei* is that it possesses a single mitochondrion per cell that is elongated along the cell body (Zikova 2022; Chaudhuri et al., 2006). Mitochondrial DNA in this organism is a concatenated structure of many circular DNAs known as kinetoplast (Jensen and Englund, 2012). In spite of the complex structure of this mitochondrial genome, it encodes only 18 proteins, the rest of the mitochondrial proteins, which is about ∼900, are nuclear-encoded and therefore imported into the mitochondrion similar to other eukaryotes (Peikert et al., 2017). However, it has been shown that mitochondrial protein import machinery in *T. brucei* is quite divergent (Chaudhuri et al., 2020; Schneider 2022, Singha et al., 2012; Harsman et al., 2016). The ATOM complex in the MOM of *T. brucei* consists of several trypanosome-specific proteins along with Atom40, the archaic homologue of Tom40 (Pusnik et al., 2011). It also performs similar functions of the TOM complex. *T. brucei* MIM possesses a single TIM complex that presumably imports presequence-containing matrix proteins as well as the polytopic MIM proteins (Harsman et al., 2016; Singha et al., 2021). The major component of the *T. brucei* TIM (TbTIM) complex is TbTim17, which is the homologue of the Tim17, Tim22, and Tim23 family of proteins in fungi and mammals. Additionally, TbTim17 is the single homologue of this family found in *T. brucei* (Singha et al., 2012; Weems et al., 2015). It associates with several unique Tim proteins to form a modular protein complex, which on a native gel separates into multiple complexes within the range of 300 to 1,100 kDa (Weems et al., 2015; Harsman et al., 2016). The predicted secondary structure of TbTim17 is very similar to fungal and mammalian Tim17/Tim22/Tim23 proteins (Weems et al., 2015). TbTim17 has four predicted transmembrane (TM) domains and the N- and C-terminal regions are hydrophilic, exposed in the IMS. Although the length of these regions and the loops between two consecutive TMs varies among different homologues in different species. Furthermore, the primary sequence of TbTim17 only shows 20 to 30% similarity to its homologs in other systems, which in contrast 50 to 60% homology among the yeast and mammalian Tim17/22/23 proteins (Singha et al., 2008). Therefore, how this single homolog in *T. brucei* performs the functions that is done by three isomeric proteins is the subject of intense study.

Similar to other nuclear encoded mitochondrial proteins TbTim17 is imported into the *T. brucei* mitochondrion and assembled in the TbTIM complex. As a polytopic MIM protein, TbTim17 does not have any predicted N-terminal targeting signal. It also has been shown that the deletion of the first 30 amino acid residues (AAs) did not hamper the import of TbTim17 into *T. brucei* mitochondrion (Weems et al., 2017). Therefore, TbTim17 must depend on internal targeting signal(s) (ITS) for its import. Multiple studies were performed to identify the ITS of *Saccharomyces cerevisiae* (Sc) Tim17 and ScTim23 (Kaldi et al., 1998, Davis et al., 1998). Results indicated that the region within the C-terminal half of these proteins, 181-222 AAs of ScTim23 and 102-158 AAs of ScTim17 are required for import and insertion into the MIM (Kaldi et al., 1998). Furthermore, these C-terminal regions possess a stretch of amino acid sequence containing few positively charged and hydroxylated residues, and this sequence was capable to import a reporter protein into the matrix (Kaldi et al., 1998). From these studies, authors speculated that this region which is in between the third and fourth TMs, is the import signal for ScTim17 and ScTim23. However, later studies revealed that the charged residues either in loop 1 or loop 3 of ScTim23 is not required for its import into mitochondria but needed for insertion of this protein into the MIM. Alternatively, a pair of TMs, particularly the first and fourth, is required for efficient import of ScTim23 (Davis et al., 1998).

Since *T. brucei* possesses a divergent import machinery and TbTim17 is assembled with non-canonical TbTims, we have systematically analyzed the structural motifs needed for import and insertion of TbTim17 into *T. brucei* MIM. Our results showed that in spite of some similarities, a few distinct features of TbTim17 are critical for import of this essential protein into the *T. brucei* mitochondrion.

## 2. Results

### 2.1. N-and C-Terminal deletion mutations revealed the importance of TM1 and TM4 for import of TbTim17 into mitochondria

It has been shown previously that the first 30 AAs, the N-terminal hydrophilic region, of TbTim17 is not required for import of this protein into the mitochondrion in *T. brucei* (Weems et al., 2017). To determine the ITS, we created a series of additional deletion mutants of TbTim17 that removed TM1 (ΔN50), TM1-TM2 (ΔN100), TM1-TM3 (ΔN120), and TM4 (ΔC31). Each mutant protein was attached to green fluorescence protein (GFP) at the C-terminal end for localization tracking in the cell (Fig. 1A). The mutant proteins were expressed in *T. brucei* from a tetracycline-inducible expression vector. The full-length (FL) TbTim17 was also attached with GFP at the C-terminal end (Fig. 1A) and expressed in *T. brucei* in a similar manner. After induction of expression, cells were harvested and sub-cellular fractions were isolated. Proteins in the total, cytosolic, and mitochondrial fractions were analyzed by TbTim17 and GFP antibodies. The cytosolic and mitochondrial marker proteins, TbPP5 and VDAC, respectively, were detected by specific antibodies, as controls. The parental T. brucei cell line (Pro 2913) were used in parallel (Fig. 1B). Results showed that the FL-TbTim17-GFP was mostly present in the mitochondrial fraction (Fig. 1C). A smaller fraction was found in the cytosolic fraction, which is likely due to overexpression of the protein. It has been shown previously that the C-terminally tagged TbTim17 (TbTim17-2X-Myc and TbTim17-tandem affinity purification, TAP, tag) were properly targeted to *T. brucei* mitochondria (Singha et al., 2012). Therefore, we do not think tagging at the C-terminal with GFP has any adverse effect on import. In contrast to the FL-TbTim17, ΔN50-TbTim17-GFP and ΔN100-TbTim17-GFP were not present in the mitochondrial fraction, but found in the cytosolic fraction (Fig. 1D & E), indicating that deletion of TM1 and TM1-TM2, hampered import of TbTim17 into mitochondria in *T. brucei*. In comparison to the FL-TbTim17-GFP, the mutant proteins were expressed less, which is likely due to degradation of the unimported proteins in the cytosol. Interestingly, ΔN120-TbTim17-GFP was primarily localized in mitochondria, showing that the C-terminal region (121-152 AAs) including loop 3, TM4, and the C-terminal hydrophilic region is capable to import GFP into mitochondria (Fig. 1F). Furthermore, when we deleted this region, the mutant protein, ΔC31-TbTim17-GFP, stayed primarily in the cytosol (Fig. 1G). We did not observe any differences in the localization of the endogenous TbTim17 in the mutant cell lines (Fig. 1B-G). Similarly, TbPP5 and VDAC were found primarily in the cytosolic and mitochondrial fractions, respectively in all cell lines, as expected. Quantitation of the immunoblot results from 3 independent experiments revealed that deletion of the first 50 or 100 AAs of TbTim17 reduced its import >98% and 100%, respectively, in comparison to the FL-TbTim17-GFP (Fig. 1H). In contrast, ΔN120-TbTim7-GFP localized to the mitochondria ∼50% in comparison to the FL-TbTim17-GFP (Fig. 1H). In addition, deletion of this region, ΔC31-TbTim17-GFP, hampered its import into mitochondria >99%. These results strongly suggest that the C-terminal 31 AAs must have some targeting information. However, it was puzzling to see that this region is present in the ΔN50-TbTim17-GFP and ΔN100-TbTim17-GFP mutants but these mutants were not imported into mitochondria, which indicates two possibilities; 1) besides the C-terminal 31 AAs, additional regions in the first two TMs are necessary for importing TbTim17, or 2) truncation of TM1 and TM1-TM2, changed the conformation of the protein so the C-terminal importing signal was not exposed. To understand, if the truncation mutations have an effect on the overall structure of the protein, we have performed in-silico structural modeling using a) RaptorX program (https://bio.tools>raptorx) with no template and b) swiss modelling (swissmodel.expasy.org) based on the Cryo-EM structure of the human Tim22 (Fig. S1A & B). From both analysis we found that the remaining overall structure of TbTim17 after truncation of TM1 and TM2 is somewhat maintained. Furthermore, similar truncations mutants of ScTim23 have been used for import studies (Davis et al., 1998). Together, our results indicate the presence of more than one targeting signals in TbTim17. As we found deletion of the first 30 AAs did not but deletion of the first 50 AAs hampered the import of TbTim17 significantly, we estimated that TM1 (31-50 AAs) likely harbors a targeting signal. Therefore, these initial results suggest the presence of possible targeting signals within at least TM1 and the C-terminal region including TM4.

**Figure 1.**
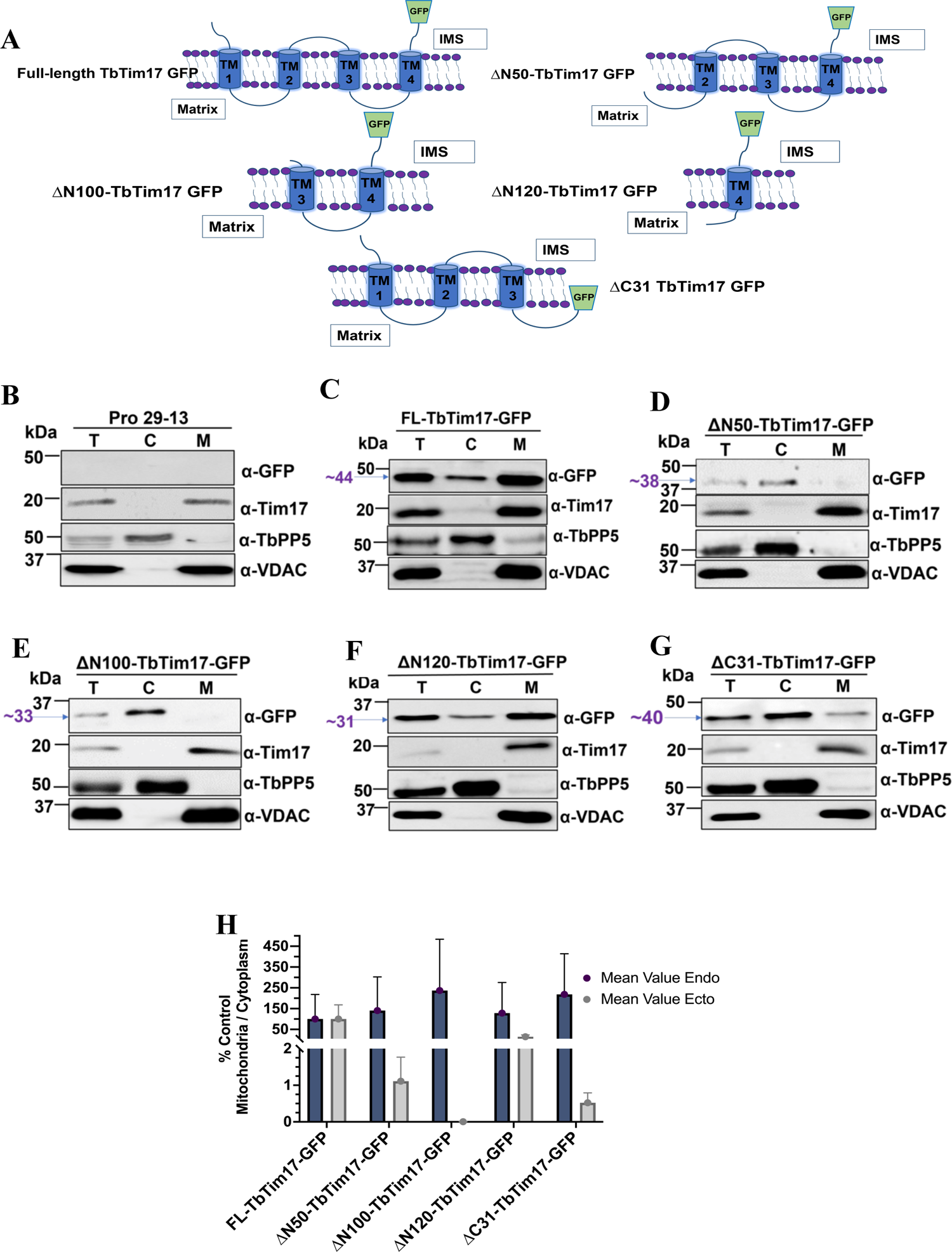
Effect of the N- and C-terminal deletion mutations on sub-cellular localization of TbTim17. (A) Schematic of N- and C-terminal deletion Mutants. The full length (FL) and mutant proteins are drawn according to the predicted membrane topology. IMS represents intermembrane space. The transmembrane domains are represented by blue cylinders and labeled 1-4. GFP tag is shown by a green trophozoite. (B-G) Immunoblot analysis of the sub-cellular fractions of the *T. brucei* expressing FL- and mutant TbTim17. The parental cell line (29-13) was used in parallel as the control (B). The stable transfectants of *T. brucei* with the FL and mutant constructs, FL-TbTim17-GFP (C), ΔN50-TbTim17-GFP (D), ΔN100-TbTim17-GFP (E), ΔN120-TbTim17-GFP (F), and ΔC31-TbTim17-GFP (G) were induced for ∼18-20 h and sub-cellular fractions were collected. Proteins in the total (T), cytosolic (C), and mitochondrial (M) fractions were analyzed by immunoblot using antibodies for GFP, TbTim17, TbPP5, and VDAC. (H) Densitometric analysis of the immunoblot results. Intensity of the GFP-fusion protein bands in the mitochondrial and cytosolic fractions were measured by Image Lab software (Bio Rad), normalized with the intensities for FL-TbTim17-GFP in the respective fractions. Densitometric analysis of the endogenous (TbTim17) in the cytosolic and mitochondrial fractions was performed in parallel. The ratio of the mitochondrial vs cytosolic fractions for both endogenous (dark blue circle) and ectopic (grey circle) were calculated as the percent of the full length protein and plotted against different cell types using GraphPad Prism. Error bars represent SEM for each data group. Sample size: n=3 (in average).

To confirm our immunoblot data we tracked the localization patterns of TbTim17 deletion mutants tagged with GFP at the C-terminal end in *T. brucei* by confocal microscopy (Fig. 2A). Mitochondria were stained with MitoTracker^TM^ Red, which is a fluorescent dye that incorporates into mitochondria in a membrane potential dependent manner (Chazotte, 2011). We induced the FL- and mutant TbTim17-GFP cell lines for ∼18-20 hours before staining with MitoTracker^TM^ Red. This shorter time period was used to avoid overexpression of the GFP-tagged protein. We used the FL-TbTim17-GFP cell line to optimize the induction period to have sufficient expression for localization of this protein then used the same time point for all other mutants. DAPI was used to stain the nuclear and mitochondrial DNAs in *T. brucei*. Afterwards, merged pictures were used to visualize co-localization of the expressed GFP-tagged proteins with MitoTracker^TM^ Red in mitochondria. We have also calculated the Pearson’s coefficient (PC) values of 15-20 individual cells for quantifying the extent of colocalization (Fig. 2B). Our results showed that the FL-Tim17-GFP protein colocalized well with the MitoTracker^TM^ Red stained mitochondria, in most of the cells as expected, with an average PC value of 0.8 (Fig. 2A & B). In contrast, we did not observe colocalization of ΔN50-TbTim17-GFP, ΔN100-TbTim17-GFP, and ΔC31-TbTim17-GFP with MitoTracker^TM^ Red as most of the GFP labelled protein was found in the cytosol of the cell (Fig. 2A). Average PC values for these three mutants were 0.58, 0.65, and 0.5, respectively (Fig. 2B). To this end, PC values near 1.0 are considered as co-localization (Dunn et al., 2011). These results were correlated with our western blot data shown in Fig. 1B. On the otherhand, ΔN120-TbTim7-GFP was colocalized with MitoTracker^TM^ Red even better than the FL-TbTim17-GFP (Fig. 2A). The average PC value for ΔN120-TbTim7-GFP was 0.95 (Fig. 2B). We noticed that ΔN50-

**Figure 2.**
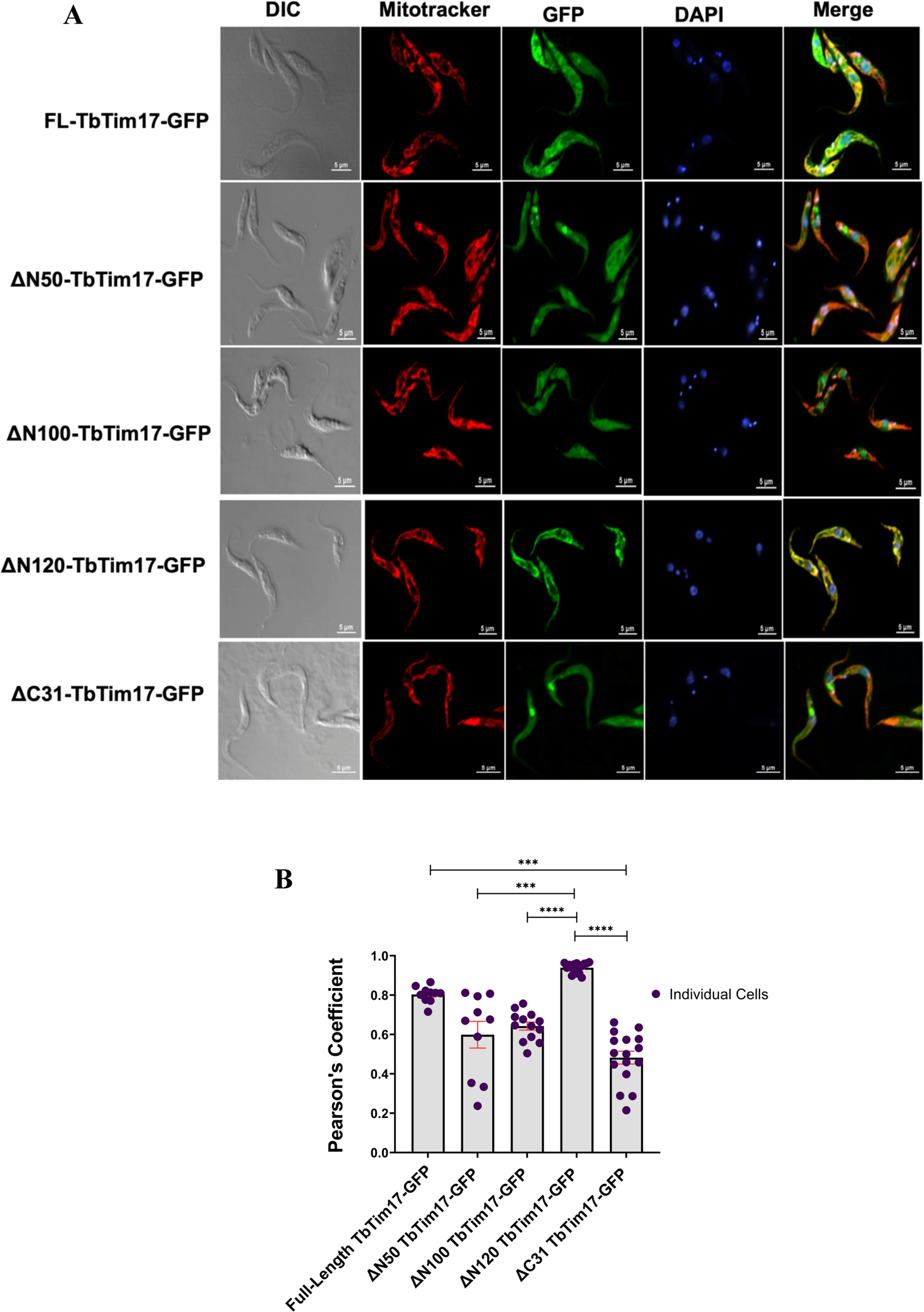
Immunofluorescence Microscopy of *T. brucei* expressed FL- and mutant TbTim17. (A) The FL-TbTim17-GFP and the mutant cell lines (ΔN50-, ΔN100-, ΔN120-, and ΔC31-TbTim17-GFP) were induced with doxycycline for 18-20 h. Cells were harvested and stained with MitoTracker^TM^ Red to visualize the mitochondrion. Expression of GFP fusion proteins are seen by green fluorescence, DAPI was used to stain the nucleus and kinetoplast (mitochondrial genome), and phase-contrast pictures (DIC) are shown. Merge images show colocalization. (B) Pearson’s coefficient values were calculated from the merge images and plotted for each type of cells using GraphPad Prism. Individual data points, n=10 (FL-Tim17 and ΔN50), n=13 (ΔN100), and n=16 (ΔN120 and ΔC31)are shown. Error bars represent SEM for each data group. Kruskal-Wallis statistical test was performed on all data sets, *** indicates p = 0.0003 (ΔN50 vs. ΔN120), p = 0.0008 (FL-17 vs. ΔC31); **** indicates p <0.0001.

TbTim17-GFP, ΔN100-TbTim17-GFP and ΔC31-TbTim17-GFP mutant proteins were accumulated in some kind of vesicles located in the cytosol of some cells (Fig. 2A). In many cases these vesicles were adjacent to the nucleus, however it requires further investigation to confirm this aberrant location of the mutant proteins. Together, our results further confirmed the importance of TM1 and TM4 of TbTim17 for its localization into mitochondria in *T. brucei*. Particulary, we found that the C-terminal region (121-152 AAs) of TbTim17 was capable of importing GFP into the mitochondria in *T. brucei*, suggesting the presence of an ITS in this region.

### 2.2. The ΔN120-TbTim17-GFP was located in the mitochondrial matrix

Next, we wanted to see if ΔN120-TbTim7-GFP is integrated into the mitochondrial membrane like the endogenous TbTim17. To investigate this, we isolated mitochondria from both FL-Tim17-GFP and ΔN120-TbTim7-GFP cell lines, treated them with 0.1M sodium carbonate at pH 11.5, and separated the soluble and pellet fractions by centrifugation. Extraction with sodium carbonate solubilizes peripherally associated membrane proteins and soluble proteins but membrane integral proteins stays in the membrane pellet (Robinson et al., 2022). After centrifugation, the supernatant and the pellet fractions obtained from both the ΔN120-TbTim7-GFP and FL-Tim17-GFP mitochondria samples were analyzed by SDS-PAGE and immunoblot analysis using GFP and Tim17 antibodies to detect both ectopic and endogenous TbTim17 proteins, respectively (Fig. 3A & B). We also used antibodies for Atom69, which is a membrane-anchored component of the ATOM complex in the outer membrane, and RNA-binding protein 16 (RBP16), which is a matrix protein. As expected, upon treatment with sodium carbonate, endogenous TbTim17 protein in both cell lines were found only in the pellet fraction indicating full integration into the mitochondrial membrane (Fig. 3A & B). Likewise, FL-Tim17-GFP protein was also found mostly in the pellet fraction although a tiny amount was released into the supernatant fraction (Fig 3A). On the other hand, a larger portion of ΔN120-TbTim7-GFP was found in the supernatant and only a tiny portion was located within the membranous pellet fraction (Fig. 3B), suggesting ΔN120-TbTim7-GFP was a soluble protein and did not integrate into the mitochondrial membrane. Therefore, although targeting GFP to mitochondria, the C-terminal hydrophilic region in conjuction with TM4 was not able to integrate the fusion protein into the membrane. It is likely that ΔN120-TbTim7-GFP is either peripherally attached to MOM and not imported into the mitochondria or it is being mis-targeted to the IMS or matrix. As expected, Atom69 was found in the pellet fractions in both cell lines, as an integral membrane protein (Fig. 3A & B). Similarly as expected, RBP16 was found primarily in the soluble fraction (Fig. 3A & B).

**Figure 3.**
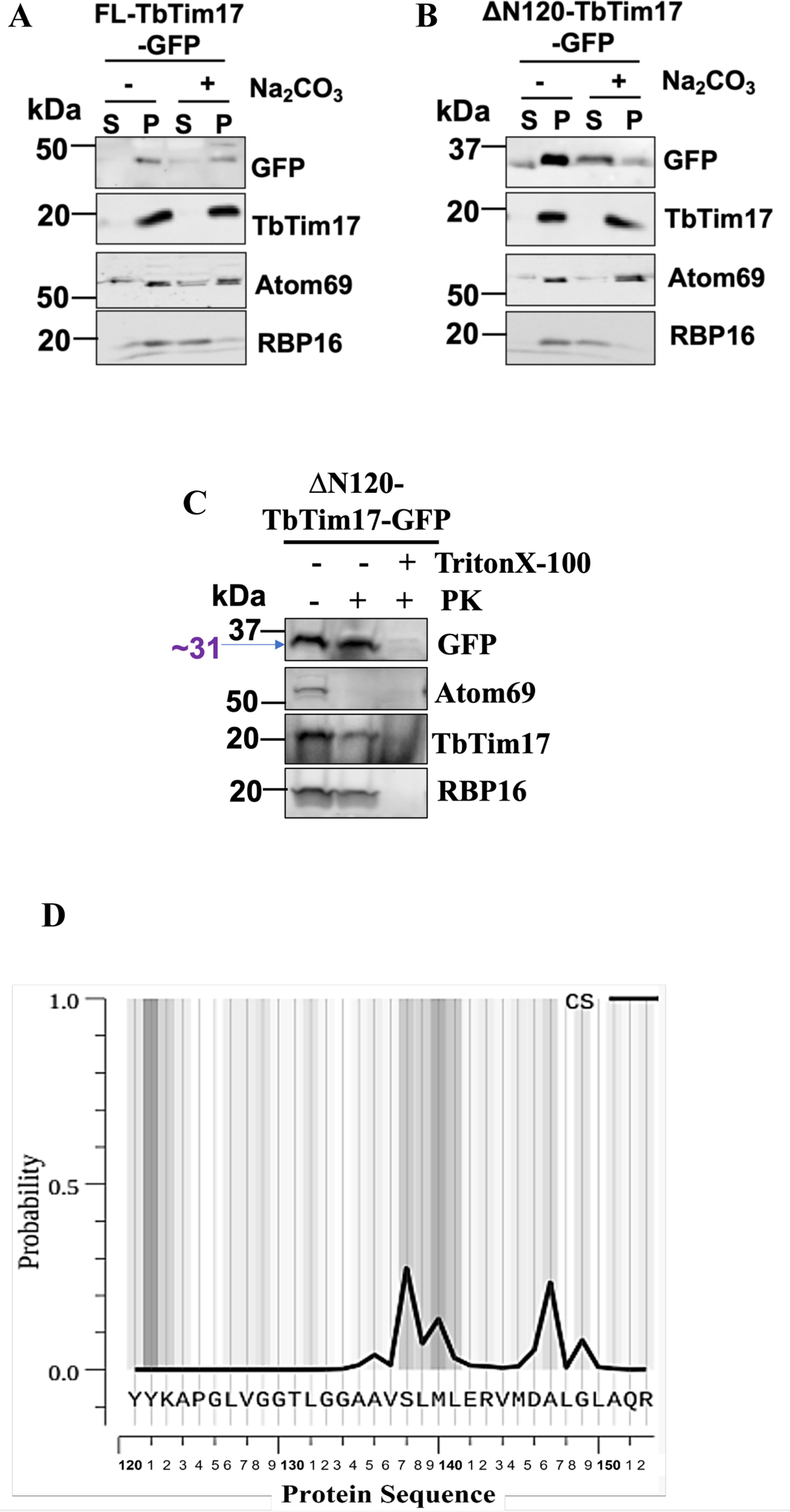
Sub-mitochondrial location of ΔN120-TbTim17-GFP. (A & B) Alkali extraction of mitochondria isolated from *T.brucei* expressed FL-TbTim17-GFP (A) and ΔN120-TbTim17-GFP (B). Mitochondria were treated (+) and left untreated (-) wth Na_2_CO_3_, pH 11.0. The soluble (S) and insoluble pellet (P) fractions were separated by centrifugation and analyzed by immunoblot analysis using antibodies as indicated. (C) Limited protease digestion of mitochondria isolated from *T. brucei* expressed ΔN120-TbTim17-GFP. Mitochondria were treated (+) or left untreated (-) with proteinase K (PK) and TritonX-100 as shown. After treatment mitochondria were recovered by centrifugation and proteins were analyzed by immunoblot using antibodies as indicated. Each experiment was repeated three times and representative blots are shown. (D) Probability plot for sub-cellular localization using TargetP 2.0 software program. The C-terminal region (120 – 152 AAs) was analyzed. The sequence of this region and the probability peaks are shown.

To investigate if ΔN120-TbTim17-GFP is peripherally associated with the MOM or it enters into the mitochondria, we treated isolated mitochondria with proteinase K (100 μg/μl). After treatment, mitochondria were pelleted by centrifugation and proteins were analyzed by western blotting using antibodies for GFP, TbTim17, Atom69, and RBP-16 (Fig. 3C). We also treated a separate portion of mitochondria with proteinase K (200 μg/μl) in the presence of Triton X-100 (1%), a non-ionic detergent, and analyzed in a similar manner. ΔN120-TbTim7-GFP was protected from digestion of proteinase K when the mitochondrial membrane was kept intact, which is similar to endogenous TbTim17 and RBP16, (Fig. 3C). In the presence of Triton X-100, proteins were digested by proteinase K, showing that the protected bands were not generated due to protease resistance. Whereas, Atom69 was completely digested by proteinase K, as expected (Fig. 3C). Together, these results showed that ΔN120-TbTim7-GFP was imported into the mitochondria, but stays as a soluble protein. As we observed that ΔN120-TbTim7-GFP was 100% protected from PK-digestion in comparison to endogenous TbTim17, which was partially (50%) digested, it is likely that ΔN120-TbTim7-GFP is located in the matrix.

It has been reported previously that ScTim17 and ScTim23, each posseses a presequence-like sequence within the TM3 and TM4. Therefore, we investigated if TbTim17 also possesses a similar signal. To determine this, we used TargetP 2.0, a software prediction program that was designed to identify subcellular localizations of proteins based on the presence of N-terminal presequences (mitochondrial, chloroplast, and Secretory/ER targeting signals) found in the primary amino acid sequence (Emanulesson et al., 2000). We analyzed residues 120-152 AAs of TbTim17 using this program and found that it has ∼41% probability for being a mitochondrial transfer peptide, ∼48% probability to act as an ER signal peptide, and ∼11% were listed as other (Fig. 3D). The primary sequence of this region consists of multiple hydrophobic residues and a few positivily charged residues (Fig. 3D). Therefore it is likely that this region being at the N-terminal of GFP acted as a presequence and translocated GFP into the matrix. Although ΔN120-TbTim7-GFP possesses TM4 of TbTim17, this TM is not strong enough to anchor the protein in the membrane.

### 2.3. TM1 alone could target GFP into mitochondria but the protein was not properly integrated into the mitochondrial membrane

As we observed that the removal of the first 30 AAs did not, but deletion of the first 50 AAs which include TM1 hampered localization of TbTim17 into the mitochondria, we decided to investigate if this region (30-50 AAs) alone could target GFP into the mitochondria. For this purpose, we created three additional protein constructs, (1) 1-30 AAs, the N-terminal hydrophilic region of TbTim17, fused with GFP [(1-30)-TbTim17-GFP)], (2) 30-50 AAs, which comprises TM1 only, linked with GFP [(30-50)-TbTim17-GFP)], and (3) 1-50 AAs of TbTim17 fused with GFP [(1-50)-TbTim17-GFP)] (Fig. 4A). These fusion proteins were expressed from an inducible expression vector as described for other mutants and sub-cellular location of these proteins were assessed by immunoblot analysis and confocal microscopy. Immunoblot analysis of the sub-cellular fractions showed that (1-30)-TbTim17-GFP was present in the cytosolic fraction (Fig. 4B), further confirming that TbTim17 does not contain an N-terminal localization signal. Conversely, the (30-50)-TbTim17-GFP localized to the mitochondria at least partially (Fig. 4C). Surprisingly, 1-50 AAs of TbTim17 failed to localize GFP to the mitochondria despite containing TM1 (Fig. 4D). Quantitation of the immunoblot results showed that about 60% of the (30-50)-TbTim17-GFP was enriched in the mitochondrial fraction, whereas enrichment of (1-30)-TbTim17-GFP and (1-50)-TbTim17-GFP was less than 5% (Fig. 4E). Together, these results indicated that TM1 indeed contains an ITS, whereas, 1-30 amino acid residues of TbTim17 somehow blocked the recognition of this signal in the (1-50)-TbTim17-GFP mutant.

**Figure 4.**
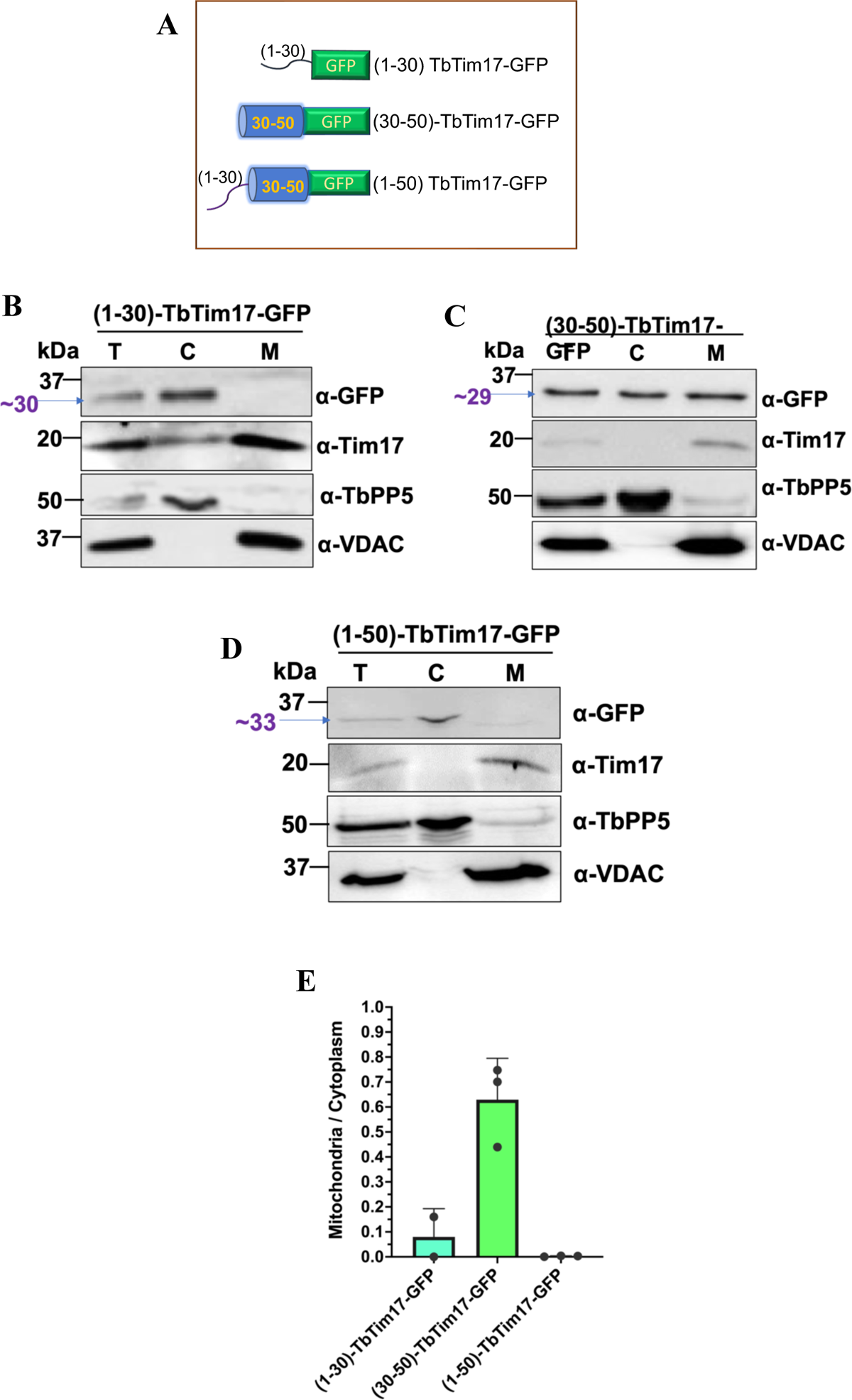
Analysis of the sub-cellular locations of the reporter protein containing TbTim17 N-terminal hydrophilic region, TM1, and both. (A) Schematics of the 1-30-, 30-50- and 1-50-TbTim17 tagged with GFP constructs. The N-terminal helical region, TM1, and GFP are shown in black line, blue cylinder, and green rectangle, respectively. (B) Immunoblot analysis of the subcellular fractions, total (T), cytosol (C), and mitochondria (M), of *T. brucei* expressing the (1-30)-TbTim17-GFP (B), (30-50)-TbTim17-GFP (C), and (1-50)-TbTim17-GFP (D) fusion proteins using antibodies for GFP, TbTim17, TbPP5, and VDAC. (E) Densitometric analysis of the immunoblot results. Intensity of the GFP-fusion protein bands in the mitochondrial and cytosolic fractions were measured by Image Lab software (Bio Rad). The ratio of the mitochondrial vs cytosolic fractions were calculated and plotted against different cell types using GraphPad Prism. Error bars represent SEM for each data group. Sample size: n=3 (in average).

We verified our western blot data with confocal microscopy (Fig. 5A). We found that the (30-50)-TbTim17-GFP colocalized with Mitotracker with an average PC value around 0.82 (Fig. 5A & B). Whereas, (1-30)-TbTim17-GFP and (1-50)-TbTim17-GFP were mostly localized in the cytosol (Fig. 5A & B), with an average PC values of 0.5 (Fig. 5B). Next we examined if (30-50)-TbTim17-GFP was integrated into the mitochondrial membrane. Alkali extraction followed by immunoblot analysis of the soluble and pellet fractions showed that a majority of the (30-50)-TbTim17-GFP present in the supernatant and a smaller fraction stays in the pellet, indicating that the protein was partially integrated into the mitochondrial membrane (Fig. 5C). Therefore, TM1 alone was not able to anchor strongly the fusion protein in the membrane. As described above we performed PK-protection assay to assess the sub-mitochondrial location of the (30-50)-TbTim17-GFP. We found that this mutant protein is well protected from PK-degestion (Fig. 5D), indicating that it crossed the MOM and partly integrated into the MIM.

**Figure 5.**
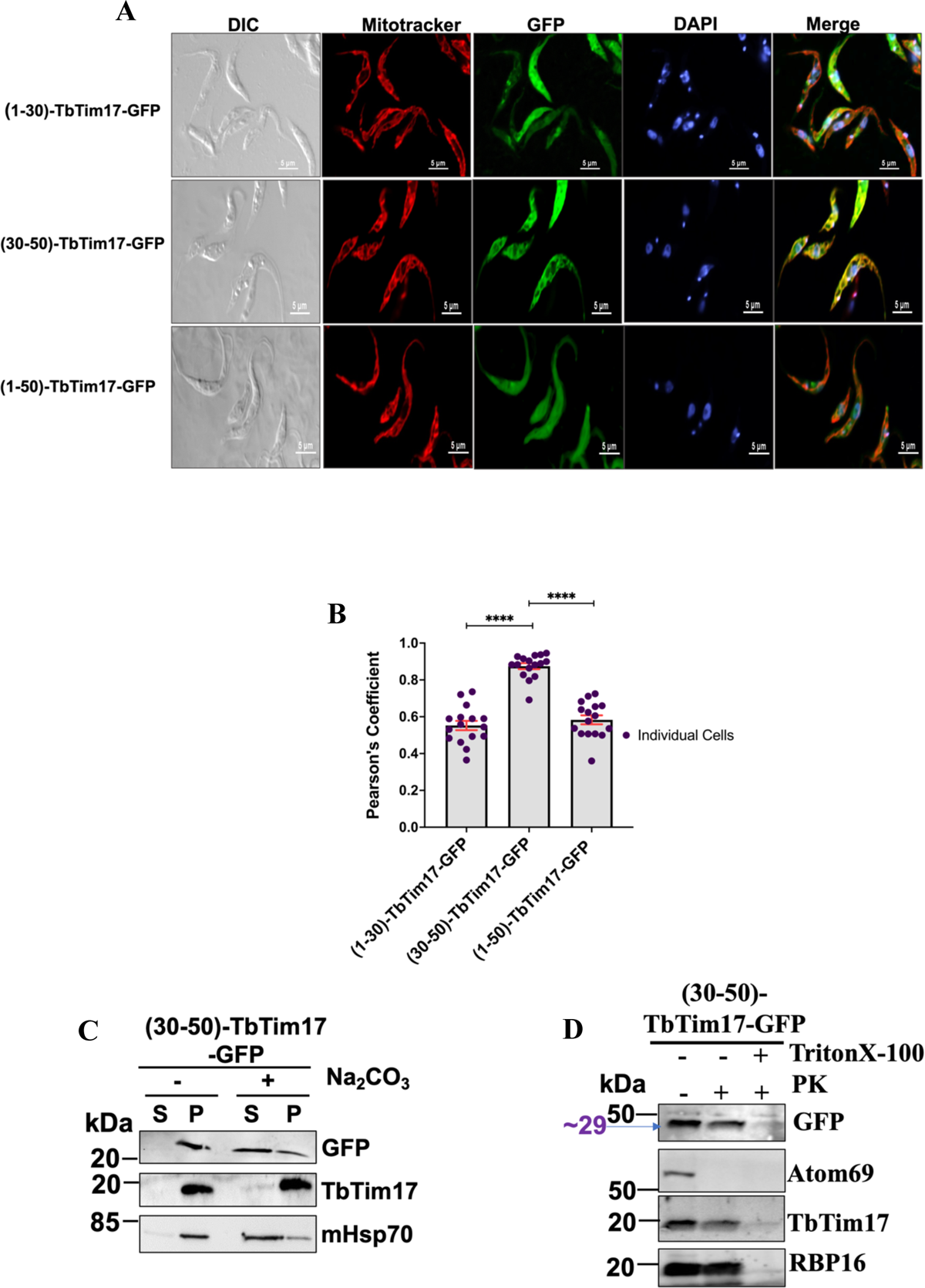
Immunofluorescence microscopy images of *T. brucei* expressed reporter protein containing TbTim17 N-terminal hydrophilic region, TM1, and both. (A) *T. brucei* cell lines, (1-30)-TbTim17-GFP, (30-50)-TbTim17-GFP, and (1-50)-TbTim17-GFP were induced with doxycycline for 18-20 h. Cells were harvested and stained with MitoTracker^TM^ Red to visualize the mitochondrion. Expression of GFP fusion proteins were seen by green fluorescence, DAPI was used to stain the nucleus and kinetoplast (mitochondrial genome), and phase-contrast pictures (DIC) are shown. Merge images show colocalization. (B) Pearson’s coefficient values were calculated from the merge images and plotted for each type of cells using GraphPad Prism. Individual data points (n=10-20) are shown. Error bars represent SEM for each data group. Statistical significance was calculated by Kruskal-Wallis statistical test, ****indicates p<0.0001. (C) Alkali extraction of mitochondria isolated from *T.brucei* expressed (30-50)-TbTim17-GFP. Mitochondria were treated (+) and left untreated (-) wth Na_2_CO_3_, pH 11.0. The soluble (S) and insoluble pellet (P) fractions were separated by centrifugation and analyzed by immunoblot analysis using antibodies as indicated. (D) Limited protease digestion of mitochondria isolated from *T. brucei* expressed (30-50)-TbTim17-GFP. Mitochondria were treated (+) or left untreated (-) with proteinase K (PK) and TritonX-100 as shown. After treatment mitochondria were recovered by centrifugation and proteins were analyzed by immunoblot using antibodies as indicated. Each experiment was repeated three times and representative blots are shown.

### 2.4. Deletion of the C-terminal hydrophilic regions and a part of TM4 did not hamper targeting but inhibits the import and integration of TbTim17 into the MIM

Since we observed the C-terminal 31 amino acid residues are required for localization of TbTim17 into the mitochondria, we wanted to investigate this region further to delineate the actual sequence responsible for mitochondrial targeting. For this purpose we generated additional mutants by deleting 2 amino acid residues at a time starting from the 10 AAs of the C-terminal end, e.i., ΔC10-TbTim17-GFP, ΔC12-TbTim17-GFP, ΔC14-TbTim17-GFP, ΔC16-TbTim7-GFP, ΔC18-TbTim17-GFP, ΔC22-TbTim17-GFP, and analyzed the localization pattern of these proteins by immunoblot analysis and confocal microscopy. Our immunoblot results of the subcellular fractions revealed that all these mutants were targeted to mitochondria, albeit at different efficiency (Fig. 6A-F). The endogenous TbTim17 and VDAC were localized in the mitochondrial fraction and TbPP5 was in the cytosolic fraction, as expected (Fig. 6A-F). Quantitation of the immunoblot results showed enrichment of the mutant proteins 2-8 folds in the mitochondrial than cytosolic fractions, except for ΔC18-TbTim17-GFP (Fig. 6G). The expression levels of ΔC18-TbTim17-GFP was also very poor, indicating the protein was unstable. Interestingly, ΔC22-TbTim17-GFP was targeted to mitochondria better than ΔC18-TbTim17-GFP, indicating that A^134^ and V^135^ at the end of ΔC18-TbTim17-GFP could be inhibitory for targeting. We also perfomed immunofluorescence localization of these proteins by confocal microscopy. We found that ΔC10-TbTim17-GFP, ΔC12-TbTim17-GFP, ΔC14-TbTim17-GFP, and ΔC16-TbTim17-GFP mutants were colocalized with Mitotracker stained mitochondria in most of the cells, which correlated with our immunoblot data (Fig. 7A). However, ΔC18-TbTim17-GFP was poorly co-localized with the mitotracker-stained mitochondria and ΔC22-TbTim17-GFP were colocalized in some of the cells, but not in others (Fig. 7A). The average PC values for the colocalization for ΔC10-TbTim17-GFP, ΔC12-TbTim17-GFP, and ΔC14-TbTim17-GFP were about 0.9, whereas that for ΔC16-TbTim17-GFP, ΔC18-TbTim17-GFP and ΔC22-TbTim17-GFP, were 0.8, 0.78, 0.8, respectively (Fig. 7B), which is slightly lower than the PC values for ΔC10 to ΔC14 mutants. These results were somewhat correlated with our immunoblot results of the subcellular fraction shown in Fig. 6, where we found a decline in mitochondrial localization of ΔC18 mutants relative to others. Overall, we found that up to a 16 AAs deletion from the C-terminus had no effect; however, more than 16 AAs deletion had a slightly negative effect on TbTim17 localization in mitochondria. We showed that complete deletion of TM4 (deletion of 31 AAs from the C-terminus) disrupts mitochondrial localization of TbTim17. Therefore, these data together indicated that residues 120 to 130 AAs likely has the critical information for mitochondrial targeting.

**Figure 6.**
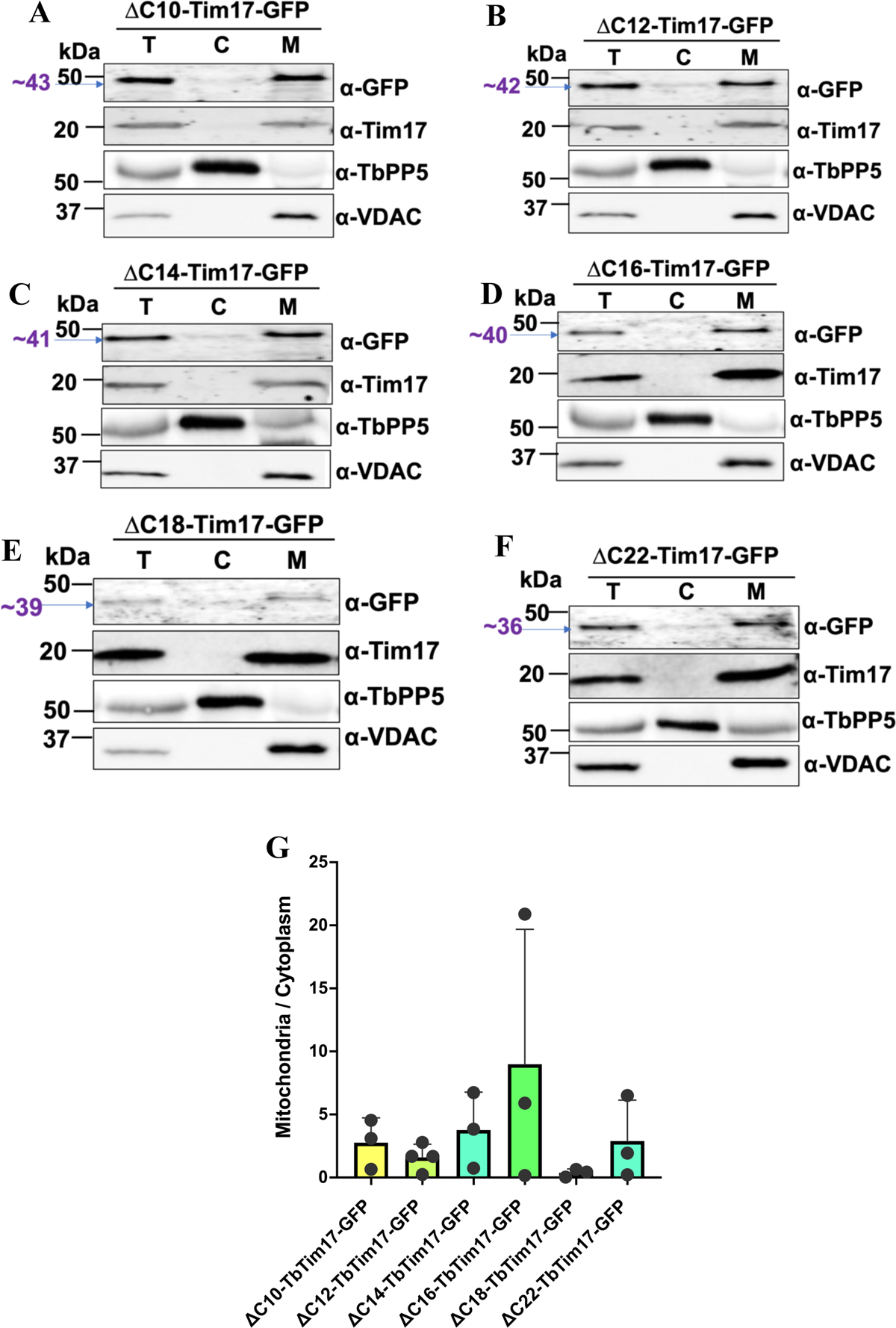
Sub-cellular location of the C-terminal deletion mutants of TbTim17. (A-F) Immunoblot analysis of the sub-cellular fractions of *T. brucei* expressed C-terminal deletion mutants of TbTim17. *T. brucei* cells expressing ΔC10-TbTim17-GFP (A), ΔC12-TbTim17-GFP (B), ΔC14-TbTim17-GFP (C), ΔC16-TbTim17-GFP (D), ΔC18-TbTim17-GFP (E), and ΔC22-TbTim17-GFP (F) were induced for 18-20 h and sub-cellular fractions were collected. Proteins in the total (T), cytosolic (C), and mitochondrial (M) fractions were analyzed by immunoblot using antibodies for GFP, TbTim17, TbPP5, and VDAC. G) Densitometric analysis of the immunoblot results. Intensity of the GFP-fusion protein bands in the mitochondrial and cytosolic fractions were measured by Image Lab software (Bio Rad). The ratio of the mitochondrial vs cytosolic fractions were calculated and plotted against different cell types using GraphPad Prism. Error bars represent SEM for each data group. Sample size: n=3 (in average).

**Figure 7.**
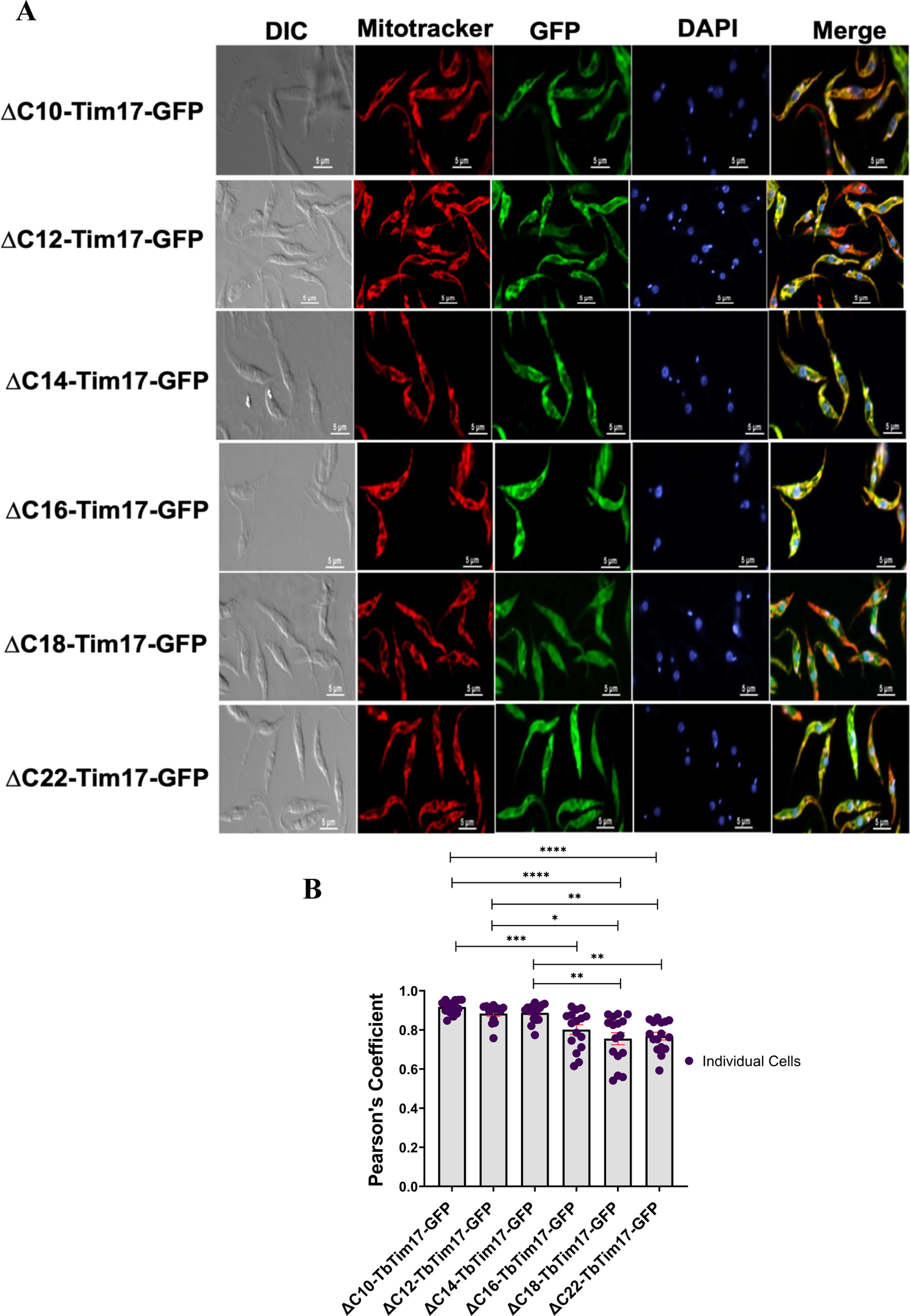
Immunofluorescence microscopy images of *T. brucei* expressing the C-terminal deletion mutants fused with GFP. (A) *T. brucei* cell lines, ΔC10-TbTim17-GFP, ΔC12-TbTim17-GFP, ΔC14-TbTim17-GFP, ΔC16-TbTim17-GFP, ΔC18-TbTim17-GFP, and ΔC22-TbTim17-GFP were induced with doxycycline for 18-20 hours. Cells were harvested and stained with MitoTracker^TM^ Red to visualize the mitochondrion. Expression of GFP fusion proteins were seen by green fluorescence, DAPI was used to stain the nucleus and kinetoplast (mitochondrial genome), and phase-contrast pictures (DIC) are shown. Merge images showecolocalization. B) Pearson’s coefficient values were calculated from the merge images and plotted for each type of cells using GraphPad Prism. Individual data points (n=10-20) are shown. Error bars represent SEM for each data group. Statistical significance was calculated by Kruskal-Wallis test, *indicates p = 0.0141; **indicates p = 0.0086 (ΔC12 vs. ΔC22); p = 0.0036 (ΔC14 vs. ΔC18); p = 0.0020 (ΔC14 vs. ΔC22), ***indicates p = 0.001, ****indicates p<0.0001.

We also wanted to see if these C-terminal deletion mutants were membrane integrated and entered into the mitochondria. Alkali extraction of the mitochondrial fractions showed that the ΔC10 to ΔC16 fusion proteins were partly present in the pellet fraction and the rest in the supernatant (Fig. 8A), indicating that the C-terminal end of the TM4 is needed for membrane integration of TbTim17. Interestingly, ΔC18-TbTim17-GFP and ΔC22-TbTim17-GFP were found mostly in the pellet fractions after alkali extraction (Fig. 8A), suggesting that deletion of the six AAs region (^131^LGGAAV^136^) relived the inhibition for membrane integration. PK-protection assay showed that except for ΔC10-TbTim17-GFP, none of these C-terminal deletion mutants were protected from protease digestion (Fig. 8B), indicating that these mutant proteins didn’t cross the MOM, possibly stuck within the ATOM channel. Therefore, it showed that TM4 of TbTim17 is required for the entry into the mitochondria. In contrast, deletion of the 10 AAs hydrophilic region of the C-terminal minimally affected the import and assembly of TbTim17 into the mitochondria. Together these data showed that the C-terminal end of the TM4 is not required for targeting but essential for translocation of TbTim17 through the MOM.

**Figure 8.**
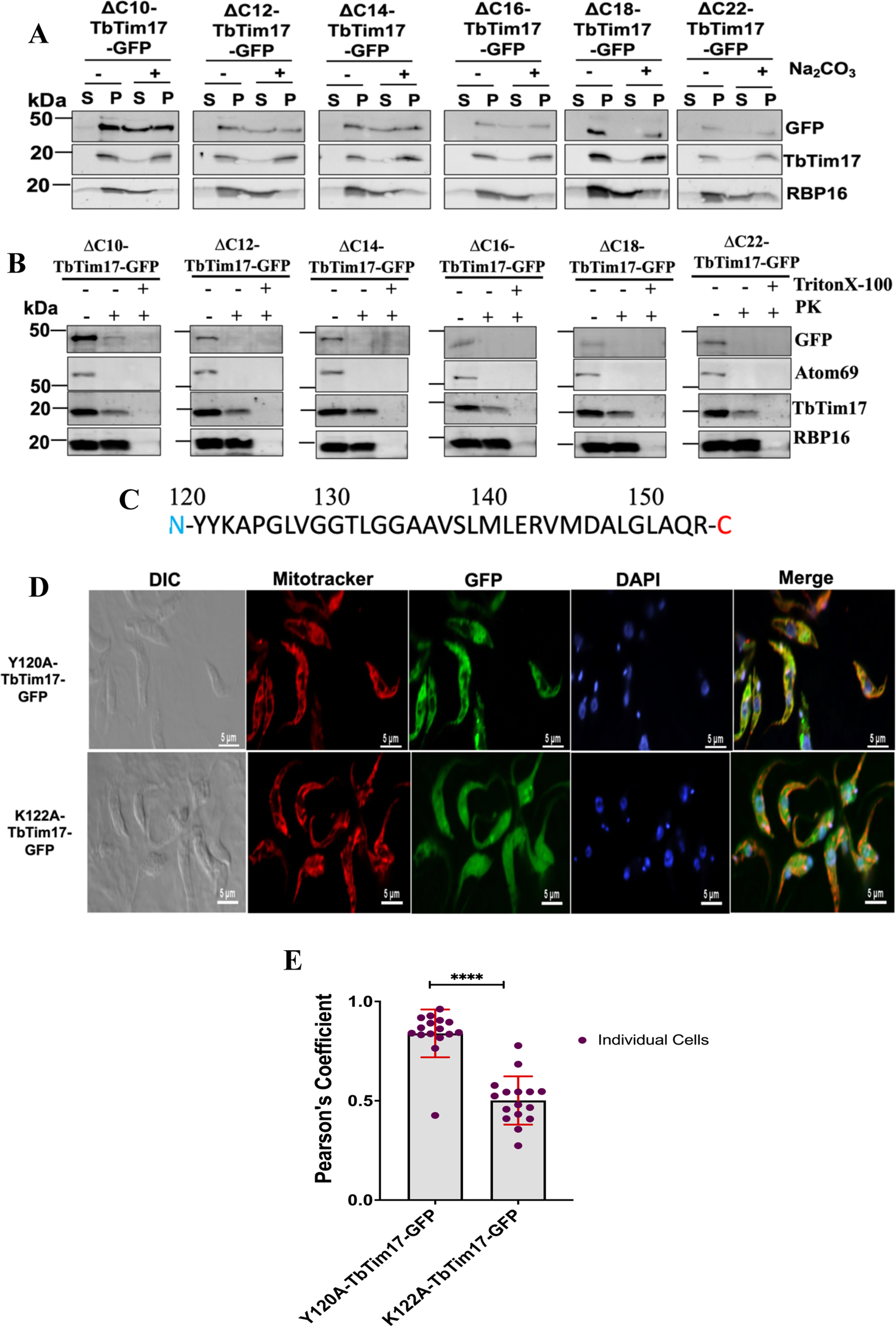
Sub-mitochondrial location of C-terminal deletion mutants and the effect of point mutation on mitochondrial localization of TbTim17. (A) Alkali extraction of mitochondria isolated from *T. brucei* expressed ΔC10-TbTim17-GFP, ΔC12-TbTim17-GFP, ΔC14-TbTim17-GFP, ΔC16-TbTim17-GFP, ΔC18-TbTim17-GFP, and ΔC22-TbTim17-GFP. Mitochondria were treated (+) and left untreated (-) wth Na_2_CO_3_, pH 11.0. The soluble (S) and insoluble pellet (P) fractions were separated by centrifugation and analyzed by immunoblot analysis using antibodies as indicated. (B) Limited protease digestion of mitochondria. Mitochondria from ΔC10-TbTim17-GFP, ΔC12-TbTim17-GFP, ΔC14-TbTim17-GFP, ΔC16-TbTim17-GFP, ΔC18-TbTim17-GFP, and ΔC22-TbTim17-GFP expressed *T. brucei* were treated (+) or left untreated (-) with proteinase K (PK) and TritonX-100 as shown. After treatment mitochondria were recovered by centrifugation and proteins were analyzed by immunoblot using antibodies as indicated. Each experiment was repeated three times and representative blots are shown. (C) Sequence of the C-terminal region of TbTim17. (D) Immunofluorescence microscopy images of *T. brucei* expressed Y120A and K122A TbTim17-GFP. Induced cells were harvested and stained with MitoTracker^TM^ Red to visualize the mitochondrion. Expression of GFP fusion proteins were seen by green fluorescence, DAPI was used to stain the nucleus and kinetoplast (mitochondrial genome), and phase-contrast pictures (DIC) are shown. Merge images show colocalization. (E) Pearson’s coefficient values were calculated from the merge images and plotted for each type of cells using GraphPad Prism. Individual data points (n=10-20) are shown. Error bars represent SEM for each data group. Statistical significance was calculated by the Wilcoxon test, ****indicates p<0.001.

### 2.5. Point mutations of Y^120^A did not, but K^122^A inhibits import of TbTim17 into mitochondria in *T. brucei*

From our observation we found that 120-130 AAs of TbTim17 (Fig. 8C) possesses information that are important for targeting and assembly of TbTim17 to mitochondria. Therefore to identify critical AAs within this region we performed site-directed mutagenesis analysis. Within this stretch of sequence, ^120^YYKAPGLVGGT^130^, there are very few polar residues like ^120^Y, ^121^Y, and ^122^K, the rest of the AAs are hydrophobic. We mutated two of these polar residues, ^120^Y and ^122^K to A individually. Mutant clones were sequenced for further verification (Fig. S2 A & B) and transfected to *T. brucei* to generate stable cell lines. Subcellular location of Y^120^A-TbTim17-GFP and K^122^A-TbTim17-GFP mutants was monitored by confocal microscopy. Results revealed that Y^120^A-TbTim17-GFP colocalized well with Mitotracker stained mitochondria; however, K^122^A-TbTim17-GFP did not show good colocalization (Fig. 8D). The average PCs were observed 0.95 and 0.55, for Y^120^A-TbTim17-GFP and K^122^A-TbTim17-GFP, respectively (Fig. 8E). Overall, our results showed that the N-terminal part of the TM4 of TbTim17 possesses an ITS, in which the single charged residue K^122^ is critical for localization of TbTim17 to mitochondria.

## 3. Discussion

We have characterized at least two ITSs for TbTim17. We showed that the ITSs are within the 1) TM1 and in the 2) C-terminal region that includes the 3^rd^ loop and TM4. Hydrophillic regions at the N- and C-termini, (1-30 AAs) and (140-152 AAs), respectively, are not required either for targeting or import of TbTim17 into the mitochondria. Both ITSs are required in combination for proper import and integration of TbTim17 into the MIM. Finally, K122, the single charged residue within the 3^rd^ loop is critical for TbTim17 import. This is the first characterization of ITS of a polytopic MIM protein in *T. brucei*.

The nuclear-encoded polytopic MIM proteins, like mitochondrial carrier family proteins (MCPs) and the mitochondrial translocase proteins, Tim17/Tim22/Tim23 family, use ITSs for their import into mitochondria (Dietmeier et al., 1993; Kaldi et al., 1998; Chacinska, 2009; Ferramosca and Zara, 2013; Horten et al., 2020; Rampelt et al., 2020). A mostly studied protein in this regard is the mitochondrial ADP/ATP carrier in *S. cerevisiae*, known as AAC/ANT (Kurz, 1999; Endres et al., 1999; Kunji et al., 2020). The AAC protein has three modular domains, each consists of a pair of TMs and a linker region facing the matrix (Endres et al., 1999; Ferramosca and Zara, 2013; Eaglesfield and Tokatlidis, 2021). Studies indicated that each module has independent targeting information but they work in cooperation to translocate AAC through the TOM complex to the TIM22 complex (Sirrenberg et al., 1996; Endres et al., 1999). Among these, the third module acts as the dominating factor for translocation of AAC (Ferramosca and Zara, 2013). On the otherhand, *in vitro* studies using isolated mitochondria and radiolabelled ScTim23 protein, Davis et al. showed that among 4 TMs, TM1 and TM4 of ScTim23 likely possess the targeting signals (Davis, 1998). However TM1 and TM4 alone were poorly imported, not protected from protease digestion and was not inseted into the MIM (Davis, 1998). The same study also showed that the positively charged residues within the connecting loops between two consecutive TMs (TM1 and TM2, and TM3 and TM4) are required for insertion of Tim23 into the MIM but not for targeting to mitochondria (Davis, 1998).

Here, we investigated the targeting signal of TbTim17 using *in vivo* studies. First we found that both TM1 and TM4 of TbTim17 have mitochondrial targeting information and these signals individually were capable to translocate a reporter protein through the ATOM channel to a protease protected location in mitochondria. However, none of these TM-containing fusion proteins were inserted into the MIM, thus remained as soluble proteins in mitochondria. Second, our studies showed that TM4 along with TM3 and the linker between them (ΔN100-TbTim17-GFP) was not targeted to mitochondria. This indicates that the presence of TM3 may hinder the recognition of the signal in TM4 of TbTim17. Similarly, we found that the TM1-GFP, [(30-50)-TbTim17-GFP)], was imported but addition of the N-terminal hydrophilic region to TM1, reduced targeting of the fusion protein, [(1-50)-TbTim17-GFP)]. This is in contrast to ScTim23, where TM1-TM2 and TM3-TM4 constructs were imported into mitochondria though at lesser extent than TM1-TM4 construct of ScTim23 (Davis, 1998). Therefore our results suggest that TbTim17 is likely imported not as loop structures as has been found for polytopic proteins in yeast. We postulated that for import and insertion of the full-length TbTim17 into the MIM, signals in TM1 and TM4 work in cooperative manner possibly via interaction with the translocase subunits. Third, we found that the C-terminal half of the TM4 in TbTim17 is dispensible for targeting but essential for its translocation through the MOM. It is likely that this region is needed to release the importing TbTim17 from the ATOM channel. Therefore, absence of this region caused the mutant proteins (ΔC12- to ΔC22-TbTim17-GFP) to be accessible to protease digestion. Thus, TM4 plays a dominant role for TbTim17 to cross the ATOM channel. Further deletion of the C-terminal region hampered the targeting and import of the mutant protein (ΔC31-TbTim17-GFP), thus the protein was found in the cytosol. Therefore, the second ITS must be within 120-136 AAs of TbTim17. Within this region, we found that the single charged residue, K122, is absolutely necessary for import of TbTim17 into the *T. brucei* mitochondrion. In contrast of having a single charged residue in loop 3 of TbTim17, the equivalent region in ScTim17 and ScTim23 have multiple positively charged residues and these were shown to be required for insertion of these proteins in the MIM (Davis et al., 1998). Therefore, the single charged residue in TbTim17 perfoms a similar job, which could be possible due to different architecture of the translocase complexes in this parasite. Together, we found that although there are some conserved features of ITSs in TbTim17 with that for ScAAC and ScTim23, the import process of TbTim17 appears distinct in *T. brucei*. *T. brucei* ATOM has two receptor subunits, Atom46 and Atom69 (Harsman and Schneider, 2017).

In our previous finding we showed that TbTim17 interacts with both of these proteins (Singha et al., 2012; Soto-Gonzalez et al., 2022), suggesting that ITSs could be recognized either by Atom46 or Atom69, or by both at the same time. Recently, we also found that each of the small TbTims (TbTim9, TbTim10, TbTim11, TbTim12, TbTim13, and TbTim8/13) is capable to directly interact with both the N- and C-terminal fragments of TbTim17, although at a greater extent with the later (Quinones-Guillen et al., 2023 preprint). In addition, we showed evidence that mitochondrial chaperone TbmHsp90/TbTRAP1 interacts with TbTim17 and is required for the assembly of the TbTIM17 complex (Soto-Gonzalez et al., 2022). Altogether, we speculate that once the ITSs of TbTim17 are recognized by the Atom receptors, the N-terminus translocate through the ATOM channel, whereas the C-terminus stays attached to the Atom subunits. Subsequently, the C-terminus is released from the ATOM channel and TbTim17 is translocated to the MIM with aids of small TbTims and by TRAP1. *T. brucei* only possesses a single TIM complex, TbTIM17. Therefore, it is likely that TbTim17 is finally translocated through this complex and assembled with other TbTim subunits via chaperons and assembly factors.

Overall, we identified the regions of TbTim17 that is crucial for TbTim17 import into mitochondria. Further investigation will identify the *T. brucei* proteins interacting with these region and will reveal the import mechanism of an essential mitochondrial protein in *T. brucei*.

## 4. Materials and Methods

### Reagents

Hygromycin, geneticin (G418), phleomycin, and blasticidine were purchased from Invivogen, minimal essential medium, restriction enzymes were from Thermofisher, Mitotracker red is from Molecular probe, aminoacids, doxycycline, buffers were from Sigma-Aldrich.

### T. brucei strain and culturing

The procyclic form of the *T. brucei* 427 double resistant cell line (29-13) expressing a tetracycline repressor gene and a T7 RNA polymerase was grown in SDM-79 medium supplemented with 10% fetal bovine serum, G418 (15 µg/ml), and hygromycin (50 µg/ml) (Wirtz et al., 1998). Cells were inoculated at 2-3 × 10^6^/ml and allowed to grow in a tissue-culture flasks at 27 ∘C incubator. Cell growth was monitored by counting cell number in a Neubauer hemocytometer under microscope.

### Plasmid constructs and transfection

To generate different deletion constructs for TbTim17, the corresponding regions of the TbTim17 open reading frame was PCR amplified with sequence-specific primers (Table S1). The forward and reverse primers were designed to add restriction sites for HindIII and XbaI at the 5ʹ ends. The PCR products were cloned in pRPX-GFP plasmid (Alsford et al., 2005) within the HindIII and XbaI restriction sites. Plasmid DNA was linearized by NotI digestion and transfected into *T. brucei* 29-13 cells, as previously described (Singha et al., 2012). Transfected cells were selected with blasticidine (10 µg/ml).

### Protein structure modeling

Three dimensional structure prediction tools, Raptor X server (Xu, 2019a; Xu, 2019b; Xu, 2020; Wang, 2017a; Wang, 2017b; and Wang, 2017c) were used to obtain predicted structures of the FL- and truncated TbTim17 mutants. Structure homology modeling of the FL- and TbTim17 mutants was performed using using Swiss Model prediction software within Chimera X based on the cryo EM structure of HsTim22.

### Sub-cellular fractionation

Fractionation of *T. brucei* procyclic form cells was performed as described (Smith et al., 2018). Briefly, 2×10^8^ cells were pelleted and re-suspended in 500 μL of SMEP buffer (250 mM sucrose, 20 mM MOPS/KOH, pH 7.4, 2 mM EDTA, 1 mM phenyl methyl sulfonyl fluoride (PMSF)) containing 0.03% digitonin and incubated on ice for 5 min. The cell suspension was then centrifuged for 5 min at 6,800 x g at 4°C. The resultant pellet was considered as the crude mitochondrial fraction, and the supernatant contained soluble cytosolic proteins.

### SDS-PAGE and Immunoblot analysis

Proteins from whole cell lysates, cytosolic, or mitochondrial extracts were separated on a 12% Tris-SDS polyacrylamide gel, transferred to nitrocellulose membrane, and immunodecorated with polyclonal antibodies for TbTim17 (Tb927.11.13920) (Singha et al., 2008), VDAC (Tb927.2.2510) (Singha et al., 2010), TbPP5 (Tb927.10.13670) (Chaudhuri, 2001), RBP16 (Tb927.11.7900) (Hayman et al., 1999), and mtHsp70 (Tb927.6.3740) (Effron et al., 1993). Antibodies for the GFP were purchased from ThermoFisher. Blots were developed with appropriate secondary antibodies and an enhanced chemiluminescence kit (Pierce).

### Alkali Extraction

For sodium carbonate extraction, mitochondria (100 μg) were incubated with 100 μL of 100 mM sodium carbonate (pH 11.5) on ice for 30 minutes. The lysate was centrifuged at 14,000 x g and the supernatant and pellet fractions were collected for further analysis.

### Proteinase K digestion

For limited proteinase K (PK) digestion, mitochondria in SME buffer at 1mg/ml concentration were treated with various concentrations of PK (0-200 ug/ml) for 30 min on ice. After incubation, PK was inhibited by PMSF (2 mM) and mitochondria were reisolated by centrifugation at 10,00 × g at 4 C for 10 min.

### Confocal Microscopy

Live *T. brucei* cells (5 × 10^6^) expressing TbTim17 mutants were used for MitoTracker staining as previously described (Smith et al., 2018). Briefly, cells were washed twice with PBS and spread evenly over gelatin-coated slides. Once the cells had settled, the slides were washed with cold PBS to remove any unattached cells. The attached cells were fixed with 3.7% paraformaldehyde and permeabilized with 0.1% Triton X-100. After blocking with 5% non-fat milk for 30 min, the slides were washed with 1X PBS. DNA was stained with 1μg/ml 4’,6-diamidino-2-phenylindole (DAPI). Cells were imaged using a Nikon TE2000E widefield microscope equipped with a 60 × 1.4 NA Plan Apo VC oil immersion objective. Images were captured using a CoolSNAP HQ2 cooled CCD camera and Nikon Elements Advanced Research software.

## Acknowledgements

We thank Sam Alsford for pRPX-GFP vector, George Cross for *T. brucei* 29-13 procyclic cells, Laurie Read for RBP16 antibody, and Paul Englund for mHsp70 antibody.

## Data availability

Datasets will be made publicly available at the time of publication.

